# Bone Resists Fatigue Through Crack Deceleration at the Fibril Scale

**DOI:** 10.1101/2025.09.30.679500

**Authors:** Riti Sharma, Stephen Ching, Luc Capaldi, Kailin Chen, Xianghui Xiao, Ottman A. Tertuliano

## Abstract

Bone endures millions of cycles throughout its lifetime by accumulating damage at a rate slow enough to allow for cell-mediated repair, but the mechanisms that delay this fatigue failure remain poorly understood. While prior studies have focused on the fatigue response of macroscale architecture of bone, the role of it’s nanoscale structure in resisting fatigue has been experimentally inaccessible. Here, we combine *in-situ* fatigue loading with synchrotron X-ray tomography and radiography to directly observe crack propagation in human bone with ∼ 21 nm spatial and 100 ms temporal resolution. We find that mineralized collagen fibrils decelerate crack growth through branching along the fibril axes, while orthogonal cracks are intermittently decelerated by nanoscale interfibrillar interfaces. These mechanisms suppress damage accumulation under physiological loads by an order of magnitude. Our findings uncover a previously unobserved toughening strategy at the nanoscale, providing insight as to how the hierarchical structure of bone bridges the timescale gap between mechanical damage and biological repair.

## Introduction

Bone is a dynamic, load-bearing tissue that persistently responds to mechanical stress by balancing damage accumulation and cell-mediated repair. The mechanics of bone have primarily been investigated under single overload conditions where it exhibits an anomalous combination of strength and toughness due to its hierarchical architecture, which can prevent^1–3^, deflect^4–8^, and even arrest fast growing fractures^6,7^. Yet, bone physiologically undergoes fatigue damage by progressively accumulating microstructural defects during repeated cyclic loading. Unlike traumatic fractures caused by a single overload event, fatigue fractures grow gradually on the scale of hours during activities like walking and running, culminating in microcracks^9^. If these microcracks accumulate and grow faster than they can be repaired, they evolve into catastrophic stress fractures. It remains unclear how fatigue crack growth in bone evolves at rates that allow for non-destructive damage accumulation, and when it transitions to catastrophic failure.

A significant challenge in understanding fatigue in bone arises from the disparate timescales over which fatigue damage initiates and cells repair the tissue. Mechanical fatigue damage accumulates rapidly, within hours, under physiological loading frequencies ranging from 0.5 to 2 Hz^10–12^, whereas biological remodeling via osteoclast-mediated resorption and subsequent osteoblast-driven bone matrix deposition occurs over several weeks^13,14^. This temporal mismatch creates a critical vulnerability window during which the structural integrity and survival of bone depends exclusively on its micro-and nanoscale architecture.

The extracellular matrix of bone is hierarchically composed of collagen fibrils, non-collagenous proteins, and nanocrystalline mineral bioapatite, a non-stoichiometric form of hydroxyapatite^15^. At the nanoscale, individual collagen molecules assemble into twisted, cross-linked structures, embedding bioapatite crystals and forming mineralized collagen fibrils ∼100 nm in diameter. These mineralized collagen fibrils serve as the fundamental building blocks that organize to form the 5 μm-sized lamellae^16^. In turn, the lamellae form the structural components of the bone architecture: the ∼100 μm osteons in cortical bone which comprise the majority of human bone mass and the similarly-sized beams and plates in trabecular or spongy bone found in vertebrae and at the end of long bones, serving as a mass-saving, topologically complex site of critical biological functions^17^.

This complex hierarchy has made it difficult to directly observe the mechanisms that initiate and drive fatigue fractures at the micro and nanoscale. Most investigations of bone fatigue focus on the macroscale, probing how cortical and trabecular structures respond to cyclic loading^18–21^ using classical fatigue parameters like Δ*K*_*I*_, defined as difference between the maximum and minimum intensity of the crack tip stress field during loading and unloading. Stable fatigue crack growth in bone has been reported over Δ*K* values spanning an order of magnitude 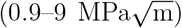, reflecting remarkable resistance to failure under repeated high intensity loading.

However, these macroscale approaches lack the spatial and temporal resolution needed to understand the underlying mechanisms at subcritical stress intensities (*<*1 MPa m^1/2^) relevant to daily physiological loading. Imaging techniques used in these macroscale studies^18,19^ have been unable to directly resolve the sub-micrometer growth rates (*<*10^−9^ m/cycle) needed to understand how fatigue evolves in a nanoscale fibrillar tissue.

Here, we introduce an experimental approach that uses synchrotron X-ray tomography and radiography combined with *in-situ* microscale fatigue loading to visualize how mineralized collagen fibrils mitigate fatigue crack growth in human bone. We quantify crack growth and arrest with 21 nm - 100 ms spatio-temporal resolution. We demonstrate fatigue toughening at the fibril level in bone by crack branching parallel to fibrils, which slows damage accumulation by an order of magnitude relative to crack propagation through fibrils. In contrast, fatigue crack propagation orthogonal to fibrils exhibits toughening by periodically decelerating when encountering interfibrillar interfaces. We find that increasing the maximum driving force during cycling loading to mimic variations in physiological loading results in a shorter fatigue life but a higher toughening rate. In short, mineralized collage fibrils and their interfaces decelerate crack propagation to prolong the fatigue life of bone and delay catastrophic failure. By revealing toughening mechanisms at the smallest structural length scale of bone, our work fills a key gap in understanding how the tissue resists fatigue damage at prior to cell-mediated remodeling.

## Results

### Fatigue in bone exhibits toughening over two crack growth regimes

Figure 1A shows a schematic of the experiment built at Beamline 18-ID, NSLS-II. Full sample and experimental preparation details are available in the Materials and Methods; briefly, human bone specimens of roughly 50 × 10 × 10 μm^3^ are fabricated using a focused ion beam (FIB) with a notch introduced as an initial flaw. The samples are manually transferred using a nano-manipulator in a scanning electron microscope onto fabricated silicon supports for the *in-situ* fatigue experiments in 3-point bending. Cyclic loading up to 10^4^ was applied using a nanoindenter equipped with a spherical tip. During loading, radiographic projection images were acquired every 100 ms to capture deformation and crack propagation dynamics. Figure 1B shows radiography snapshots of a fatigued bone specimen at 500, 6500 and 9700 loading cycles. At 500 cycles, no evident crack propagation was observed, and the bone specimen appears undeformed. By 6500 cycles, notch opening increases before any significant visible crack extension, although signs of crack branches appear near the notch tip. By ∼10^4^ cycles of fatigue loading, the radiography image shows ∼4 μm of stable and slightly tortuous crack growth.

**Figure 1:**
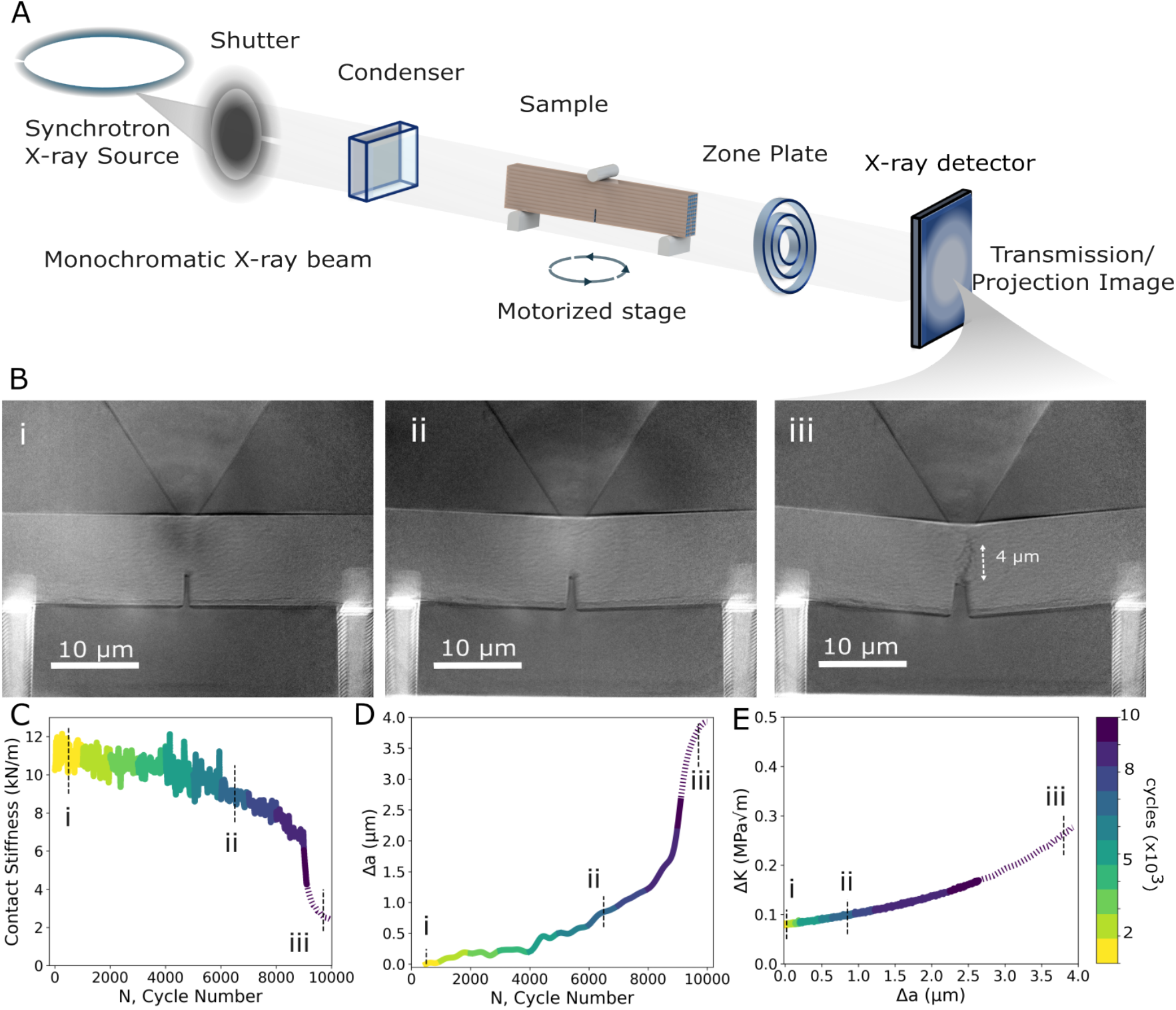
Fatigue toughening revealed via *in-situ* synchrotron X-ray loading. (A) Schematic of the *in-situ* X-ray synchrotron experiment, with specimen in beam path for radiographic imaging during fatigue. (B) Radiographic images showing crack progression in the microscale beam at i. 500 cycles, ii. 6500 cycles, and iii. 9700 cycles, highlighting notch opening and crack growth over 10^4^ cycles. (C) Evolution of contact stiffness over 10^4^ cycles. (D) Effective crack length derived from stiffness data. (E) Rising R-curve of the beam, indicating crack-growth resistance and toughening behavior. Dashed lines indicate potential crack tip interactions with the indenter stress field (see Fig. S2).

To characterize damage evolution, we tracked the stiffness of the bone specimens in Fig. 1C to calculate crack extension, Δ*a* with respect to fatigue cycle in Fig 1D. This calculated Δ*a* of 4 μm is consistent with the final crack extension measured from the radiography of Fig. 1Biii, confirming that specimen stiffness provides an accurate metric for tortuous crack extension in this nanofibril microstructure. During progressive fatigue loading, we observed a stiffness drop of up to 75% as the crack propagated. Two distinct regimes emerged from these measurements. The initial regime exhibits a progressive but steady drop in stiffness up to 8600 cycles, consistent with crack initiation followed by stable propagation. The slow decrease in stiffness suggests the activation of fatigue mechanisms that slow fatigue prior to catastrophic fracture. Beyond 8600 cycles, a second regime was observed, marked by a significantly accelerated drop in stiffness, indicative of unstable crack growth.

To understand how the slow progressive crack extension may relate to fatigue toughening, we calculated the energy dissipated during each increment of crack growth (see Supplemental). In order to propagate crack growth, the fatigue driving force (Δ*K*) had to increase. This increase results in a rising R-curve (Fig. 1E), indicating active toughening during fatigue loading. The observed R-curve can be distinctly divided into two toughening regimes consistent with our crack growth rate analysis. The initial regime, extending up to cycle 8600, corresponds to approximately 1.6 μm of crack extension and is characterized by lower crack growth rates (∼0.4 nm/cycle). Toughening then transitions to the second regime with higher crack growth rates (∼4 nm/cycle). These distinct regimes suggest a transition in underlying mechanisms that govern toughening throughout fatigue crack propagation in bone.

### Crack branching along fibrils delays initial fatigue crack progression

To investigate the toughening mechanisms that give rise to fatigue toughening, we analyzed the full crack path by comparing high-resolution 3D tomography datasets (Fig. 2A,B), acquired before and after fatigue loading. The tomographic reconstructions were obtained with a voxel size of 21 nm, enabling resolution of individual mineralized collagen fibrils within the microscale bone specimens (Fig. 2A, inset). The pre-fatigue tomography in Fig. 2A shows fibrils oriented primarily orthogonal to the notch while Fig. 2B shows the same specimen after 10^4^ cycles of fatigue loading. This deformed configuration reveals that the specimen accommodates applied stresses through both permanent deformation in bending and crack propagation from the initial notch.

**Figure 2:**
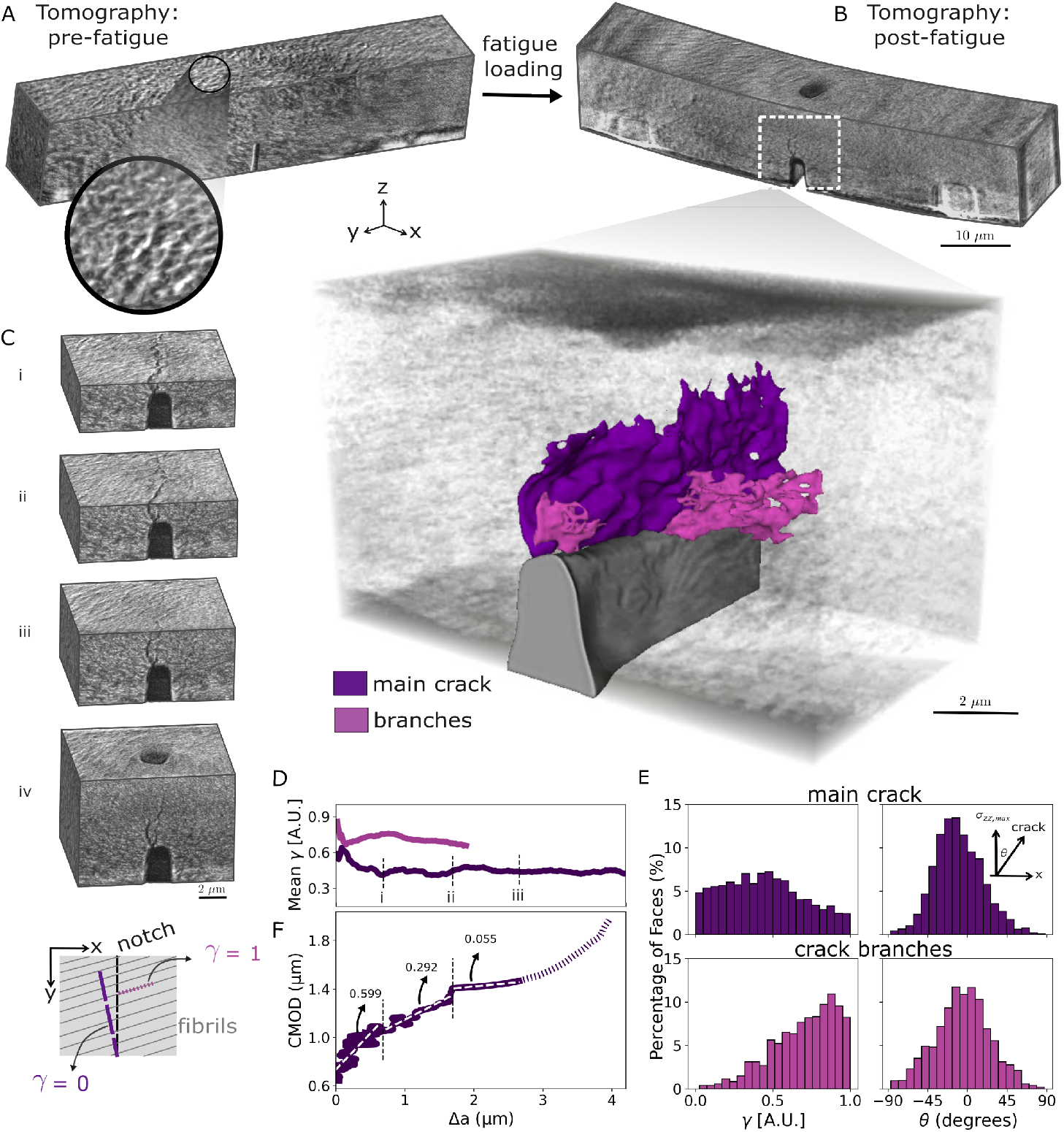
Tomography reveals fibril-level crack branching. (A) Tomographic reconstruction of the microscale bone beam prior to fatigue loading. Inset demonstrates resolving individual ∼100 nm mineralized collagen fibrils from absorption contrast. (B) 3D tomography of the same beam post-fatigue, revealing fibril deformation and crack propagation. The inset shows crack segmentation, distinguishing the main crack from secondary branches. (C) Cropped 3D tomography images at increasing crack extension (Δ*a*) intervals, showing the evolution of damage and the surrounding fibril deformations in 3D. (D) Crack alignment to fibrils (*γ*) as a function of crack length, reveals that relative to the main crack, branches exhibit higher alignment with the fibril structure. The main crack exhibits high initial alignment that decreases as the crack propagates. (E) Crack propagation angle relative to the principal tensile stress direction. Both main and branch cracks generally align with the expected path of maximum tensile stress (*θ* = 0), but branches show greater alignment with the underlying fibril structure. Over time, both converge toward the longitudinal (maximum tensile) direction. (F) CMOD data plot with derived effective crack length shows three different slope regions during the experiment; highlighting fundamentally different mechanisms in play through the extension of the crack.

To visualize how this crack propagation leads to toughening, we traced the full 3D crack path (Fig. 2B, inset) and segmented into two components: a main crack and additional branches extending from it. The main crack spans the out-of-plane thickness of the specimen and grows as a continuation of the notch, whereas branches are defined as cracks growing obliquely to the notch. Figure 2C shows cross-sections at various slices along the main direction of crack extension. At early stages of crack extension (Fig. 2Ci), the main crack has sawtooth-like fluctuations along the thickness of the specimen that appear to result from fracture of individual fibrils. These small fluctuations become less apparent at larger crack extensions as the main crack front sharpens and appears discontinuous through the thickness of the beam (Fig. 2Ciii). Although fibrils appear to introduce fluctuations, the overall morphology and orientation of the main crack (Fig. 2B) is consistent with the expected crack extension towards the direction of global maximum tensile stress in Mode-I crack opening^22^, rather than conforming to the local microstructure. Yet, the branches extending obliquely from the main crack suggest there may be a more significant contribution of the underlying fibril microstructure to fatigue toughening.

To assess this hypothesis, we examined whether the local fibril orientation near the propagating crack could explain the emergence of branches. We quantified the alignment, *γ*, of the crack faces to the local fibril orientation. We defined *γ* = 1 − |**n** *·* **f** |, where **n** is the unit normal vector that locally describes crack face orientation (Fig. S3) and **f** is the unit vector parallel to the local the fibril orientation. *γ* = 1 indicates alignment between the crack face and local fibrils, while *γ* = 0 indicate the crack is growing orthogonally to the local fibril orientation. Figure 2D shows the mean *γ* value per 21 nm thick slice as a function of crack extension, Δ*a*. The branches exhibit consistently higher alignment with the fibrils compared to the main crack throughout the fatigue process. However, during the first ∼150 nm of fatigue crack propagation, the main and branched cracks exhibit similar alignment to the underlying fibril orientation, suggesting fatigue initiation may be triggered by a nanoscale crack growth along the direction of fiber orientation. Within 700 nm of crack growth, the main crack redirects and stabilizes along the direction of maximum tensile stress. In contrast, the branches show increasing alignment with the fibrils at small crack extensions Δ*a <* 700 nm, followed by a slight decrease that still preserves this relative alignment. In Fig. 2E, we complement *γ* by quantifying *θ*, the angle between the local crack surface and the loading direction. We observe that *θ* for the main and branched cracks are normally distributed and centered around *θ* = 0 indicating that Mode-I fatigue fracture behavior drives global crack growth. Yet, the distribution of *γ* suggests a distinct role of branching, correlated to fibril orientation, in mitigating crack growth during fatigue initiation.

To assess whether branch growth plays a prominent role as a toughening mechanism during the early stages of fatigue crack growth, we plot the crack mouth opening displacement, CMOD, directly extracted from radiography images versus the stiffness-derived crack extension, Δ*a* (Fig. 2D). We postulate that branches growing transverse to the direction of loading will lead to increases in CMOD without substantial increases in Δ*a*, whereas the main cracks should increase both. In Fig. 2D, we identified three regimes based on the slope of CMOD vs. Δ*a* (Fig. 2D) that reflects the relative contribution of the branching and map to the observed changes in *γ*. The first and steepest regime, i, has a pronounced increase in CMOD relative to minimal changes in Δ*a*. This regime occurs over Δ*a <*700 nm, consistent with when branched cracks exhibit increased alignment to fibrils in Fig. 2D. The second regime, ii, occurs over 700 nm *<* Δ*a <* 1.6 μm and exhibits a decreased slope relative to regime-i. This regime represents a transition into stable re-orientation of the main and branched cracks to constant values of *γ* in Fig. 2D. The final regime, iii, shows a pronounced increase in crack extension with respect to CMOD, suggesting a transition towards catastrophic crack growth. This regime is associated with crack propagation dominated by the main crack with a lack of visible branches at Δ*a >* 1.6 μm (Fig. 1B,D). These observations suggest that branching along the direction of fibrils likely serves as a prominent toughening mechanism early in non-destructive fatigue crack growth by slowing crack extension. These results motivate an examination of how individual fibrils specifically accommodate damage during fatigue.

### Nanoscale interfaces decelerate fatigue crack growth between fibrils

Given that cracks that aligned parallel to fibril orientation delay fatigue progression by serving as branches, we asked how fatigue crack propagation occurs orthogonally to fibrils. Figure 3A shows meandering crack propagation towards the direction of loading. Here, meandering refers to small, localized deflections of the crack path at fibrillar interfaces, followed by re-alignment toward the overall loading direction. By analyzing the crack growth rate, *da/dN*, as a function of crack extension (Fig. 3B), we observed periodic oscillations in the growth rate at every 70–200 nm of crack growth. These oscillations persisted up to 1.6 μm of crack growth and suggest microstructural interactions that cause periodic deceleration of fatigue crack growth, unrelated of the loading frequency (Fig. S5).

**Figure 3:**
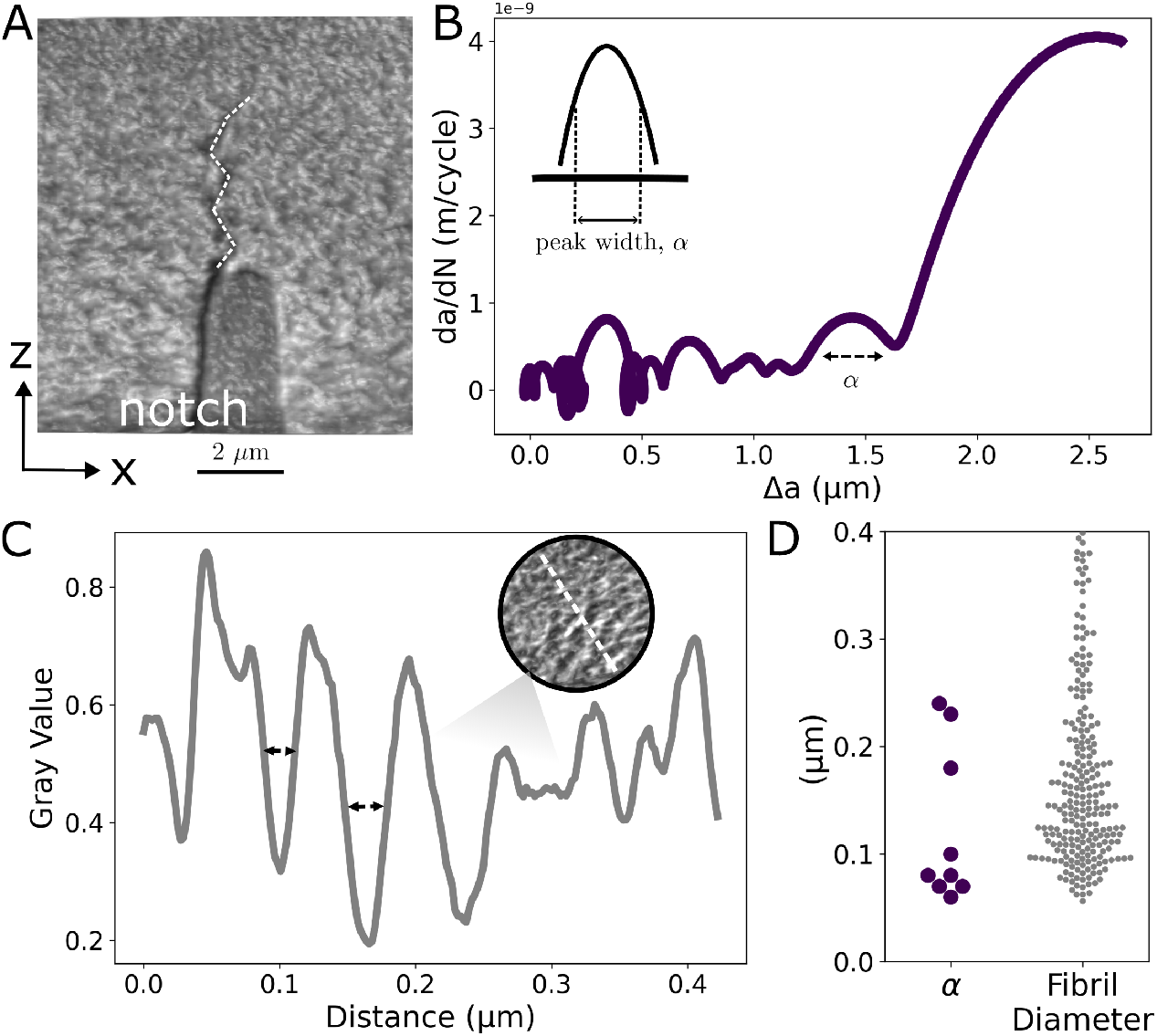
Nanoscale interfaces enable periodic fatigue deceleration. (A) Crack propagation profile in the z–x plane showing the meandering nature of crack growth. (B) Crack growth rate, *da/dN*, plotted against crack extension, Δ*a*. Periodic drops in growth rate, observed at discrete intervals between approximately 70–200 nm, reflect repeated crack deceleration events. (C) Representative intensity profile from a single slice showing mineralized collagen fibrils (bright regions) and interfibrillar spaces (dark regions). Valley widths are calculated from these oscillations as marked by the arrows. (D) Plot comparing the widths of periodic drops in crack growth rate (*da/dN*), α and the valley widths from the intensity profile. This analysis correlates the alignment of periodic crack growth rate reductions with fibrillar dimensions.

We hypothesize that these decelerations are caused by the softer interfibrillar “glue” between mineralized collagen fibrils in bone^23,24^. To support this hypothesis, we measured the periodicity of fibrils in our specimens. Figure 3C shows a representative intensity profile on a tomographically reconstructed slice of a pre-fatigue specimen. The periodic oscillations in the intensity profile, measured as the valley widths, correspond to transitions from acceleration to deceleration during crack propagation. Figure 3D shows that the distribution of the full width at half maximum (FWHM) for the peaks in crack growth rate matches closely with the periodicity observed in the fibril intensity profile. These results support that crack deceleration occurs at fibril interfaces.

This interpretation is supported by prior studies on layered materials^25,26^, which demonstrate that elastic mismatch at interfaces significantly influences crack behavior. Brach et al. demonstrated that interfaces with notable elastic mismatches influence crack propagation due to local mixed-mode loading conditions, especially under Mode-I loading, by inducing shear stresses at the crack tip. The crack naturally seeks a path of minimal shear stress, thereby deflecting along compliant interfaces to satisfy the principle of local symmetry and reduce crack-tip stress intensities^25^.

In bone, compliant extrafibrillar non-collagenous proteins (NCPs) with a modulus of approximately 1 GPa^27^ interface with stiffer mineralized collagen fibrils with a higher modulus ranging from approximately 2 *−* 5 GPa^28^. This elastic mismatch, exceeding a factor of two, promotes local deflection and decelerates crack growth. Similar phenomena have been observed in macroscale fracture studies of cortical bone, where microcracks encountering osteonal cement lines oriented perpendicular to the principal crack path act as effective delamination barriers through the Cook-Gordon mechanism^7,26^. This mechanism describes how weak interfaces blunt propagating cracks, induce significant crack meandering, and create highly tortuous crack paths as seen in Fig. 3A and Fig. 2C. Such deflections drastically reduce local stress intensities at the crack tip, requiring increased applied energy to resume crack propagation and thus significantly enhancing toughness. The observed meandering exhibit self-similarity across multiple length scales, highlighting how hierarchical elastic mismatches at the nanoscale mirror toughening strategies observed at the osteon level.

Collectively, crack branching and deceleration at fibrillar interfaces reveal a coordinated toughening strategy that operates exclusively during fatigue. Both mechanisms are active at early and slow stages of crack extension. Under cyclic loading, the gradual accumulation of damage allows the crack front to engage more directly with the hierarchical microstructure. Branched cracks dissipate energy by propagation parallel to fibrils, while crack growth deceleration at interfaces delays orthogonal crack growth. These mechanisms have not been observed under monotonic loading conditions at this scale where the driving force for fracture and crack propagation increases too rapidly for such microstructural interactions to be effective^4,29–31^.

After approximately 1.6 μm of crack extension, we observed a notable transition in the deformation behavior, characterized by significantly higher crack growth rates from approximately 0.4 nm/cycle to as high as 4 nm/cycle. At this stage, the previously dominant toughening mechanisms of crack branching and deceleration at fibrillar interfaces become largely exhausted. The subsequent overload regime has been well documented in existing literature, particularly highlighting collagen fibril bridging as a primary toughening mechanism^4,5,32,33^. In bone, fibril bridging emerges as the crack transitions from a fatigue-driven, slow-growth stage to a regime characterized by rapid propagation. This fibril bridging can be visualized in Supplemental Video 1 of the post-fatigue specimen.

This distinction highlights an important temporal sequence in bone’s fatigue resistance mechanisms. Early-stage mechanisms such as branching and interfacial deceleration function preemptively, effectively delaying failure onset, whereas fibril bridging represents a traditional shielding mechanism activated under longer cracks and elevated stress intensity conditions. Together, these findings demonstrate how bone’s multi-stage toughening response is intricately governed by its unique fibril architecture and nanoscale interfaces. This response under constant amplitude cyclic loading raises a broader physiological question: how do variations in fatigue loading affect bone’s resistance to crack propagation?

### Higher stress intensities accelerate damage but improve toughening rate

Bone is naturally subjected to cyclic loads during daily activities such as walking, running, or jumping—each imposing varying intensities that correspond to different fractions of its intrinsic fracture toughness, 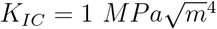. To contextualize these physiological loading conditions, we introduce a non-dimensional 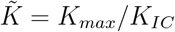, representing the maximum applied stress intensity normalized by the critical fracture toughness of bone. From a materials perspective, it is important to note that the maximum stress intensity, *K*_*max*_, plays a dominant role in brittle and quasi-brittle systems^34^, where cyclic loading drives crack growth similarly to static loading intensified by degradation of crack-tip shielding mechanisms. In our experiments, we investigate fatigue behavior under 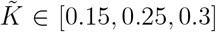, reflecting conditions ranging from regular activity to elevated loads associated with an increased risk of acute injury^35^(Fig. 4).

**Figure 4:**
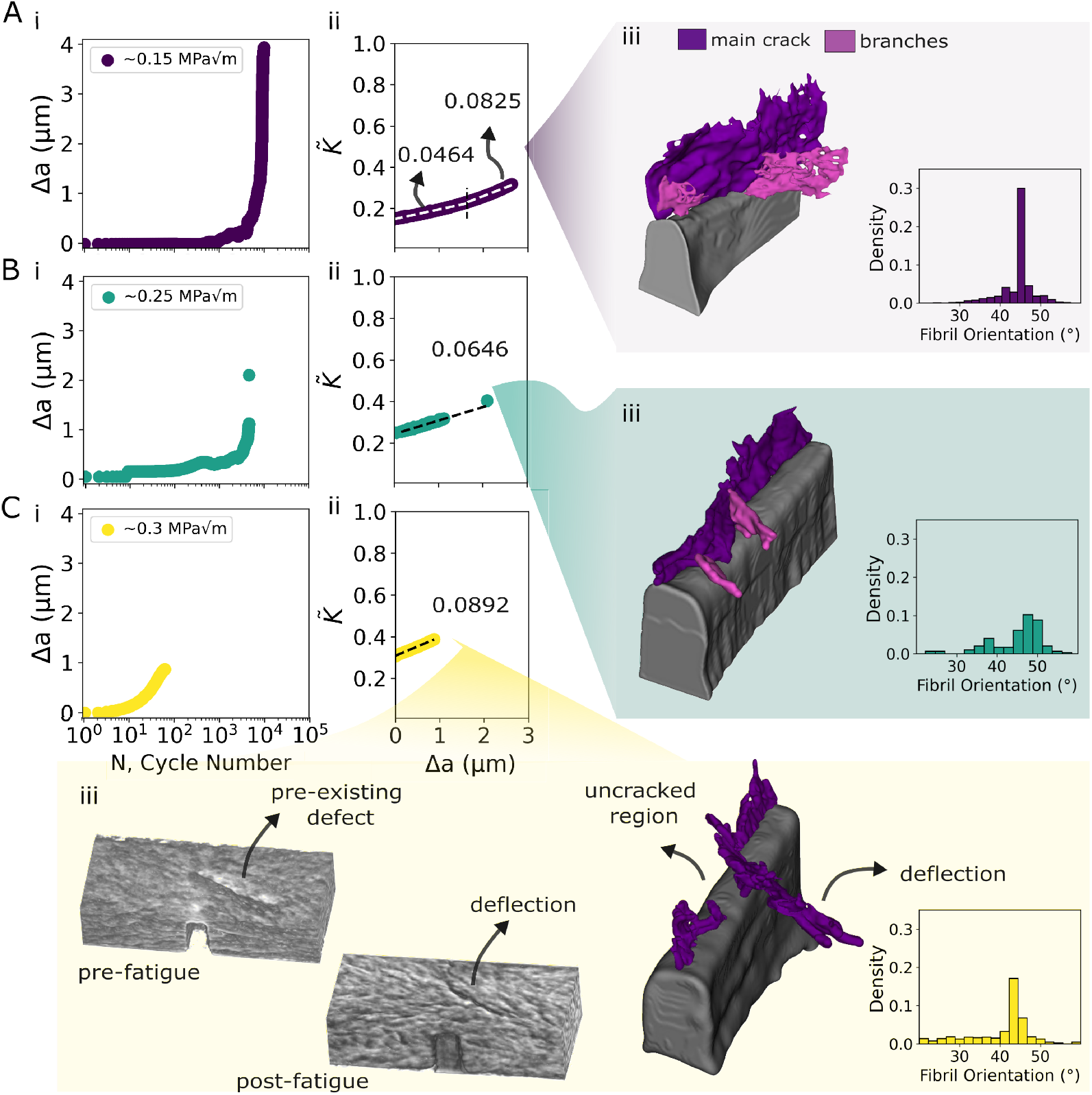
Increasing stress intensity increases toughening rate. (A) At 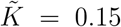, the crack extends 4 μm over 10^4^ cycles; the toughening slope doubles at the transition from fatigue to overload regime; 3D segmentation shows 4 branches during fatigue loading. (B) At 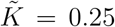, the crack extends 2.1 μm over 4700 cycles; the toughening slope steepens; and the 3D segmentation highlights the presence of two branches. (C) At 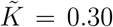, the crack propagates fastest to extend 1 μm within 65 cycles; the toughening slope is highest; 3D analysis shows a pre-existing defect forces the crack to deflect into a mixed-mode path.

The detailed fatigue toughness behavior observed in the previous sections were at 15% of the fracture toughness of bone (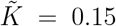, 10,000 cycles, Fig. 4Ai). This specimen distinctly highlights two toughening regimes, characterized by their differing slopes on the rising R-curve (Fig. 4Aii). In the fatigue-dominated region extending up to cycle 8600 (approximately 1.6 μm of crack extension, reaching 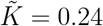), a toughening slope of 0.0464 is recorded. This regime correlates with the extensive crack branching and deceleration at fibril interfaces observed experimentally, which slowed crack growth under cyclic loading. Subsequently, transitioning into the overload regime (cycles 8600–9100), the toughening slope doubles to 0.0825, indicating enhanced fracture resistance associated with higher 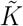 values. This regime is predominantly characterized by fibril bridging, where crack growth is impeded by bridging ligaments in the crack wake. Such a transition in dominant toughening mechanisms highlights the importance of stress intensity in activating a range of toughening mechanisms. Segmentation analysis (Fig. 4Aiii) shows the propagation of main crack and four branches due to the applied cyclic loads.

These results align with prior literature on brittle materials such as alumina (Al_2_O_3_) and silicon carbide (SiC). Short fatigue cracks in these materials show lower fatigue resistance due to limited crack bridging in their wake regions. As bridging zones develop further with increased crack length, fatigue resistance improves^36^. Drawing parallels to our experiments in bone, achieving higher 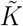 values likely promotes activation of toughening mechanisms such as crack bridging, enhancing fracture resistance beyond mechanisms dominant at lower stress intensities.

Next, by initially fatiguing at 25% of the fracture toughness of bone, (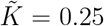, 4,700 cycles), Fig. 4Bi) we observed a shorter fatigue region up to 1.1 μm before catastrophically jumping to 2.1 μm (Fig. S5). Here, the R-curve had a toughening slope of 0.0646, an increase compared to the 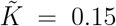 condition (Fig. 4Bii). Figure 4Biii shows the segmentation of the evolved crack, including the formation of two branches. We also observed similar deceleration of crack growth rates in the fatigue regime consistent with fatigue at lower value of 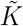 (Fig. S5). These observations suggest that while similar fatigue mechanisms are active across different loading intensities, the extent of toughening they provide is modulated by the applied 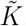.

By fatiguing at 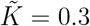, (∼65 cycles), the specimen exhibited highest crack growth rates, associated with increased susceptibility to fracture (Fig. 4Ci). Despite the high 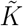 and shorter fatigue duration, this condition yielded the highest toughening slope of 0.0892, (Fig. 4Cii), approaching physiological overload conditions. Figure 4Ciii shows evidence of a pre-existing defect near the notch which influenced the crack trajectory, attracting the nucleated fatigue crack into a highly mixed-mode crack growth, and leaving an uncracked region behind. The observed crack path and deformation highlights that increasing 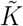 reduces the number of cycles until failure-defined here as ∼1 μm of stable crack extension-while simultaneously enhancing the rate of toughening.

These findings demonstrate higher loading intensities may shorten fatigue life but they also accelerate toughening because the bone microstructural promotes crack deflection and bridging to enhance resistance against rapid crack growth. At physiological loading frequencies of 1 Hz, common daily activities impose cyclic loads varying in magnitude and duration; a half-marathon subjects bone to approximately 10^4^ loading cycles, whereas recreational jogging involves ∼ 10^3^ cycles, and a short sprint corresponds to about 10^2^ cycles. We observed that under lower loading intensities, toughening through crack branching and deceleration at interfaces dominates to slow early fatigue damage, yielding steadier and incremental propagation. Activities involving lower-intensity loading may favor damage-stabilizing mechanisms that extend fatigue life, while higher-intensity activities could activate more aggressive toughening responses that resist single overload by accelerating the activation of fatigue mechanisms.

## Summary and Outlook

The 3D nanoscale organization of mineralized collage fibrils plays a multifunctional role in bone; it guides tissue development and resists mechanical damage. The ability to quantitatively analyze the 3D organization and nanoscale damage mechanisms within mineralized collagen fibrils is essential to understanding the structure-function relationships that dictate bone’s remarkable damage tolerance. While recent studies have emphasized the role of nanoscale collagen architecture in bone development^37^, our work reveals that these same structures are equally critical for suppressing crack propagation at the earliest stages of fatigue damage.

Leveraging a 21 nm-resolution 3D synchrotron imaging platform that surpasses previous capabilities^38,39^, our multimodal experiments directly track fatigue-toughening mechanisms *in-situ*. We find that mechanical loading intensity modulates distinct, complementary toughening strategies: higher loads preferentially promote crack deflection and fibril bridging, enhancing immediate crack growth resistance, while lower loads favor incremental crack branching and deceleration at interfaces, enabling sustained energy dissipation. These observations highlight the dynamic coupling between mechanical loading and microstructural architecture in governing fracture resistance.

Our findings provide a broader mechanistic insight: mineralized collagen organization, and specifically its interfaces, serves as a natural design motif optimized to promote both regenerative remodeling and mechanical resilience. Understanding how these roles are balanced could inform the development of architected and engineered living materials that integrate long-term structural integrity with the adaptability promised by biological responsiveness.

## Methods

### Sample fabrication

Bulk human trabecular bone samples from a 69-year-old female donor with no history of bone disease were purchased via a tissue harvesting consortium (Articular Engineering, Inc.). Samples were cleaned by successive washes in isopropanol (IPA) and deionized (DI) water, polished to a precision of 1 μm, and coated with a thin (∼ 75 *nm*) conductive Pt–Pd layer to mitigate charging effects. Microscale (50×10×10 μm^3^) beams were fabricated using the Focused Ion Beam – Scanning Electron Microscope (FIB-SEM) at Drexel University, operating at 30 keV accelerating voltage. The beams were oriented such that lamella were predominantly parallel to the beam span, replicating physiologically relevant crack-arrest orientations observed in bone. Each beam included a small notch as an initial flaw. These beams were subsequently extracted via lift-out and mounted onto silicon supports (2×2 mm^2^ wafers) using ion beam-induced platinum deposition glue, forming a microscale three-point bending geometry (Fig.1B).

### *In-situ* loading setup and imaging

Samples were transported to the Brookhaven National Laboratory (BNL) synchrotron facility for *in-situ* X-ray imaging at beamline 18-ID, employing monochromatic X-rays at 9.4 keV and obtaining a resolution of 21 nm. Nanomechanical fatigue experiments were conducted using an *in-situ* testing apparatus (KLA NanoFlip) equipped with a spherical diamond tip. The microscale beams were initially loaded at a normalized stress intensity 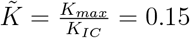, followed by a constant amplitude cyclic loading (1 Hz, R=0.5) to drive fatigue crack propagation. Mechanical parameters including load, displacement, stiffness, and cycle number were continuously recorded throughout testing. Concurrently, radiographic images were captured every 100 ms to visualize real-time spatial and temporal changes in mineralized collagen fibril structure near the crack tip. High-resolution 3D tomographic scans (21 nm resolution) were acquired before and after fatigue loading to quantify structural alterations.

### Crack length calculation

The effective crack length, Δ*a*, was calculated using the contact stiffness measured during the duration of the experiment. Using a compliance calibration procedure, the stiffness at the start of the experiment (cycle 1) was identified as the maximum measured stiffness, and inverted to determine the bone specimen’s initial compliance. The main assumption is that the minimum of the measured dynamic compliance represents the initial effective compliance, *C*_0_, of the fabricated specimen with an initial crack length *a*_0_.

The relationship between the load line compliance, *C*, and the normalized crack length, 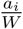, for a three-point bend fracture specimen was calculated using^40^:

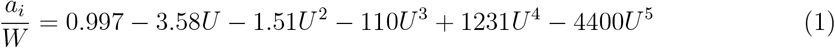

where

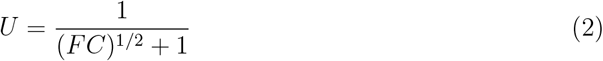

Here, *F* is a calibration constant specific to each specimen, determined using initial geometry and compliance conditions This is done by setting 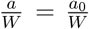 and *C* = *C*_0_ for each specimen. The instantaneous crack extension is then obtained by solving for 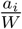 using the measured compliance *C*_*i*_. This process allows for continuous tracking of crack propagation as the specimen is subjected to cyclic loading, providing crack extension data over the entire experimental duration. This smoothed crack extension data was used to calculate crack growth rates, *da/dN*, and stress intensity factor values, 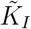 (see Supplemental).

### Radiography and Tomography processing

Radiography images were acquired during fatigue loading at a temporal resolution of 100 ms per frame. To ensure uniformity across images, flat-field and background correction was applied using the following normalization formula: 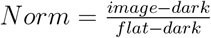. Corrected radiographs were used to calculate the crack mouth opening displacement (CMOD), as shown in Fig. S4. A video of the loading sequence, compiled from these corrected radiographs, is provided in the Supplementary Materials. Further details on CMOD extraction and analysis are also included in the Supplementary (Fig. S4).

Pre-and post-fatigue X-ray tomography was performed on microscale beams to capture the 3D arrangement of mineralized collagen fibrils and crack propagation relative to the introduced notch. Tomographic datasets were reconstructed and visualized using open-source software packages including TXM Sandbox^41,42^, Dragonfly^43^, and 3D Slicer^44^.

### Crack Segmentation

Crack paths were segmented from reconstructed tomography data using intensity thresholding in 3D Slicer^44^, leveraging contrast differences between cracks and surrounding bone matrix. Threshold values were iteratively optimized to isolate cracks accurately while minimizing artifacts. Resulting binary masks were converted to fitted 2D surface meshes along the crack’s main vertical axis (aligned with maximum tensile stress direction) and any oblique crack branches. These fitted surfaces were used to quantify crack path orientation. Cracks identified in the pre-fatigue tomography were labeled as pre-existing. By overlaying pre-and post-fatigue 3D reconstructions, these pre-existing cracks were traced in the post-fatigue dataset. Newly formed cracks were categorized either as the main crack or as crack branches. The main crack was defined as the dominant fracture extending along the notch axis, while branches were defined as cracks propagating obliquely from the notch in the xz-plane.

### Fibril Orientation Measurements

Reconstructed tomography volumes were subdivided into smaller volumes of size 2.5 μm × 2.5 μm × 0.2 μm. For each subvolume, an intensity profile was extracted by sweeping a horizontal line through the center of the medial xy-plane, rotating it incrementally through a range of angles (Fig.3C). For each rotation angle, the number of local maxima in the intensity profile was counted using a minimum peak width of 50 nm and a minimum peak prominence of 0.1. The angle that produced the greatest number of peaks was taken to be perpendicular to the in-plane collagen fibril orientation. In cases where multiple angles produced the same number of peaks, the angle with the smallest mean peak width was selected. After determining the fibril orientation, half prominence valley widths in the intensity profile were measured to characterize the size of low-intensity interfibrillar material where crack growth rate accelerates.

To evaluate crack deflection relative to fibril organization, the angle between the crack face normal and fibril orientation was quantified. Specifically, the crack-fibril alignment metric, *γ*, was defined as *γ* = 1 *−* |**n** · **f** |, where **n** is the unit normal vector of a crack face (extracted from fitted 2D mesh facets) and **f** is the unit vector representing fibril orientation in the nearest subvolume. Similarly, the crack orientation relative to the applied loading direction was defined as *θ* = 1 *−* |**n** · **e**_*y*_|, where **e**_*y*_ is the unit vector along the loading axis.

## Acknowledgments

This work is supported by OAT’s NSF CAREER Award CMMI - 2339836. This research used resources at beamline 18-ID of the National Synchrotron Light Source II, a U.S. Department of Energy (DOE) Office of Science User Facility operated for the DOE Office of Science by Brookhaven National Laboratory under Contract No. DE-SC0012704. FIB-SEM fabrication was performed using instruments in the Materials Characterization Core (RRID: SCR 022684) at Drexel University. This work was carried out in part at the Singh Center for Nanotechnology, which is supported by the NSF National Nanotechnology Coordinated Infrastructure Program under grant NNCI-2025608.

## Author contributions

OAT conceived the idea. RS, LNC, KC, and OAT performed the experiments. RS and SC analyzed the data. XX helped develop the synchrotron setup and provided guidance on image processing. RS, SC, OAT wrote, revised, and edited the manuscript. All authors discussed the results and final interpretations. OAT supervised the project.

