## Supplemental Video 2 for "Bone Resists Fatigue Through Crack Deceleration at the Fibril Scale"

### Supplemental Information

#### Correcting experimentally measured stiffness for contact compliance

The spherical indenter's contact point with the bone beam exhibits finite stiffness, contributing to the experimentally measured stiffness. To account for this, the experimental system is modeled as a series of springs, as illustrated in Fig. S1A. The measured stiffness is given by:

$$S_{\text{measured}} = \left(S_{\text{specimen}}^{-1} + S_{\text{spherical tip}}^{-1}\right)^{-1} \tag{3}$$

where $S_{\text{specimen}}$ and $S_{\text{spherical tip}}$ represent the stiffness contributions from the bone beam specimen and the spherical indenter tip, respectively. To quantify the contribution of the spherical tip, we use contact mechanics models that provide theoretical contact stiffness values based on the tip geometry. This contribution of the spherical tip, $S_{\text{spherical tip}}$ is modeled as a contact between a sphere and a flat plane. An elastic sphere of radius $R$ indents an elastic half-space where total deformation is $d$, causing a contact area of radius:

$$a = \sqrt{Rd}$$

The applied force $F$ is related to the displacement $d$ by:

$$F = \frac{4}{3}E^* R^{\frac{1}{2}} d^{\frac{3}{2}}$$

According to Hertzian contact mechanics, the stiffness of the spherical tip can be computed using the following equation:

$$S_{\text{spherical tip}} = 2E^*\sqrt{R}\sqrt{d}, \tag{4}$$

The reduced modulus $E^*$ accounts for the combined elastic properties of both the indenter and the specimen, considering their respective moduli and Poisson's ratios:

$$E^* = \left( \frac{1-\nu_1^2}{E_1} + \frac{1-\nu_2^2}{E_2} \right)^{-1} \tag{5}$$

where $E_i$ and $\nu_i$ are elastic moduli and Poisson's ratio. $E_1$ and $\nu_1$ are 1190 GPa and 0.06 for diamond, and $E_2$ and $\nu_2$ are 15 GPa and 0.3 for bone.

For this study, the indentation depth was manually measured from radiographic images at multiple cycle numbers (Fig. S1B). Red markers indicate the manually extracted indentation depths at discrete cycles, while the blue lines represent interpolated values used to estimate continuous indentation depth across all cycles (Fig. S1C). These interpolated depths were then used to compute the theoretical contact stiffness as a function of indentation depth, and then correlated to cycle numbers. These corrections reveal a stiffening in the elastic response of the specimen (Fig. S1D). This depth-based stiffness correction was applied uniformly across all specimens in this study, and the corrected data was used to compute the relevant fracture results.

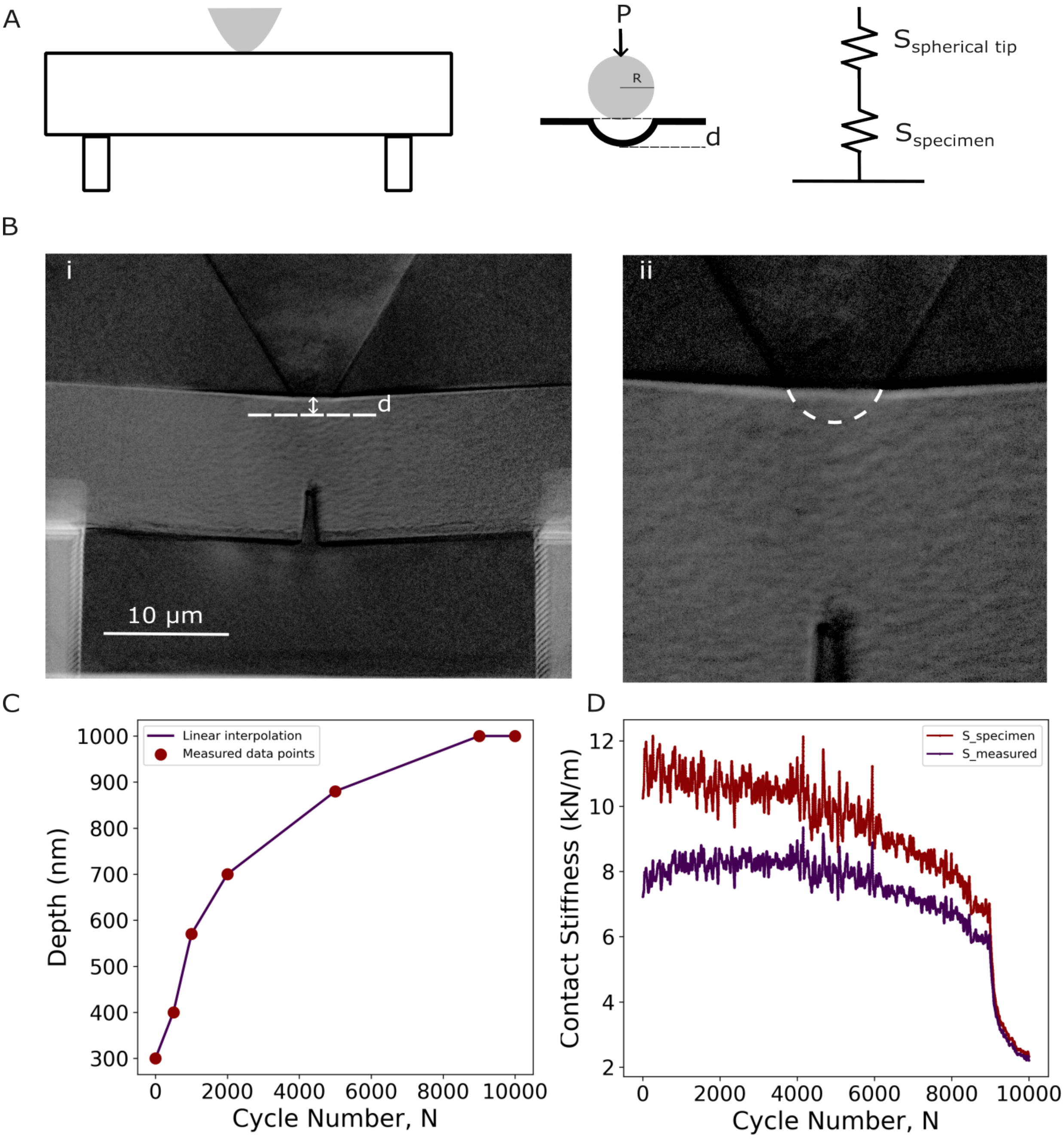


**Figure S1:** (A) Schematic of the three-point bending setup used in the experiment, along with a representation of the spherical indenter in contact with a deforming half-space to a depth, d. The system is modeled as two springs in series: the spherical contact stiffness ($S_{spherical}$) and the specimen stiffness ($S_{specimen}$). (B) Representative radiographic image showing the spherical tip indentation into the specimen. Manual measurements of indentation depth were taken at selected cycle numbers. (C) Plot of indentation depth as a function of cycle number. Red points indicate manually measured depths, while the blue lines represent interpolated values between these measurements. (D) Comparison of the original experimentally measured stiffness and the corrected stiffness after accounting for contact deformation. The correction reveals stiffening with respect to original measured values.

#### Calculating stress intensity factor, $K_I$

The mode I stress intensities, $K_{max}$, $K_{min}$, and $\Delta K$ are calculated from the load data and crack extension at each cycle using the Linear Elastic Fracture Mechanics (LEFM) relation:

$$K_I(i) = \frac{P_i S}{B W^{3/2}} f\left(\frac{a_i}{W}\right) \tag{6}$$

Here, $P_i$ and $a_i$ are the instantaneous load and crack length, respectively, and $S$ is the span of the beam. $B$ is the The term $f\left(\frac{a_i}{W}\right)$ accounts for the specific fracture geometry of the specimen. For a macroscale single-edge notched beam (SENB), this term is defined as[1]:

$$f\left(\frac{a_i}{W}\right) = \frac{3\left(\frac{a_i}{W}\right)^{1/2}\left[1.99 - \left(\frac{a_i}{W}\right)\left(1 - \left(\frac{a_i}{W}\right)\left(2.15 - 3.39\left(\frac{a_i}{W}\right) + 2.7\left(\frac{a_i}{W}\right)^3\right)\right)\right]}{2\left(1 + 2\left(\frac{a_i}{W}\right)\right)\left(1 - \left(\frac{a_i}{W}\right)\right)^{3/2}} \tag{7}$$

#### Crack tip interactions with the indenter stress field

To assess whether the indenter influenced crack propagation near the end of the fatigue test, we examined both the radiographic and tomographic datasets for signs of interaction between the crack tip and the stress field of the spherical indenter.

Figure S2A shows radiographic snapshots of the specimen and has the the overall region of interest towards the end of the fatigue loading. Figure S2B shows a zoomed-in view revealing the crack tip approaching and interacting with the indenter field. Stiffness-based analysis of crack growth in Fig. S2C reveals a sharp drop in the crack growth rate at 9100 cycles, corresponding to approximately 2.7 μm of total crack extension. This sudden decrease suggests that the crack tip has entered the stress field or plastically deformed zone associated with the spherical indenter, which likely acts as a barrier to further propagation.

Based on this observation, we define 9100 cycles as the onset of interaction with the indenter stress field. Data beyond this point is represented using dashed lines in subsequent plots, indicating that the crack behavior is no longer governed purely by the material response but may be influenced by contact interactions with the indenter.

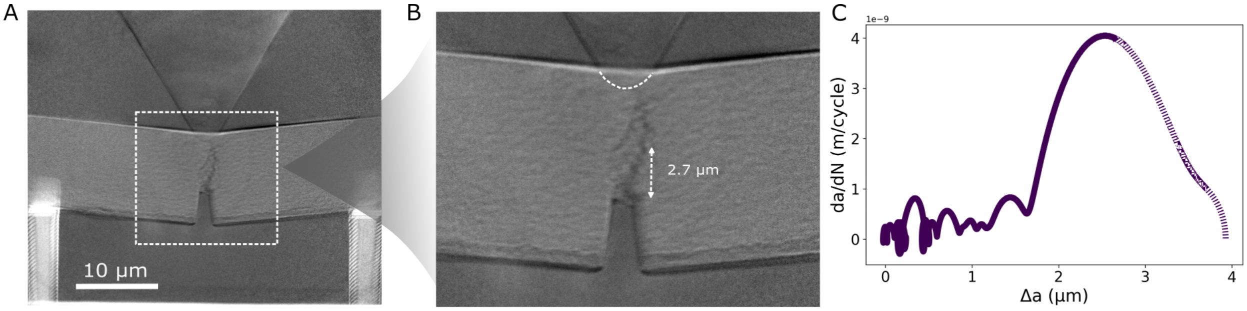


**Figure S2:** (A) Radiographic images of the specimen at the end of fatigue loading. Overview of the region of interest showing the crack approaching the indenter. (B) Zoomed-in view revealing interaction between the crack tip and the indenter's stress field. (C) Stiffness-based crack growth rate analysis showing a drop at ∼9100 cycles (∼2.7 μm )

#### Unit normal definition for segmented crack

To describe the orientation of the crack faces, we computed the unit normal vector at each element of the segmented crack surface. Figure S3 shows the segmented crack surface with the overlaid mesh used for normal vector computation. The main crack is shown in purple, while crack branches are highlighted in pink. The mesh itself is shown in black. A zoomed-in view of a representative region is provided to depict an individual mesh element with its corresponding unit normal vector. This vector locally defines the crack face orientation and is calculated for every mesh element within the segmented surface.

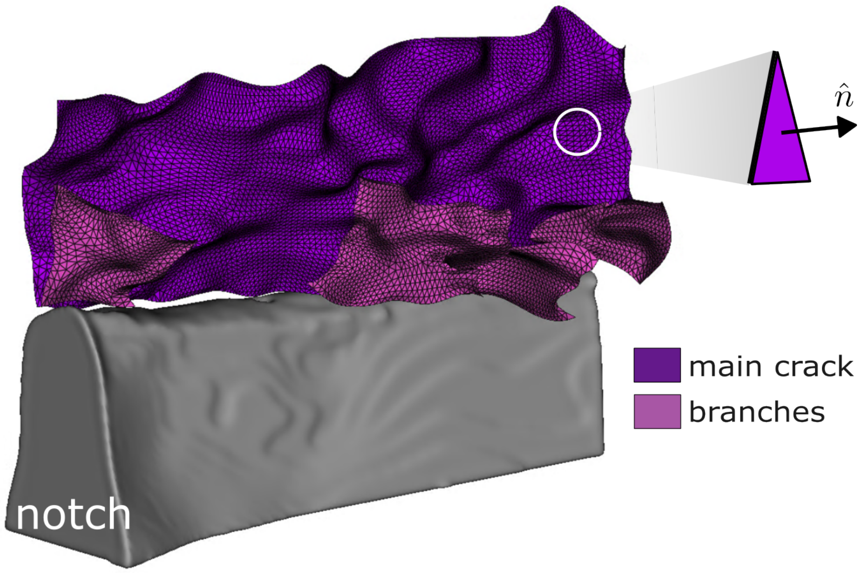


**Figure S3:** Visualization of the segmented crack surface with overlaid mesh used for unit normal vector computation. The main crack is displayed in purple, and crack branches are shown in pink. The magnified view highlights a representative mesh element with its associated unit normal vector, demonstrating how the local crack face orientation is defined throughout the surface.

#### Measurement and evolution of Crack Mouth Opening Displacement (CMOD) during fatigue loading

Crack mouth opening displacement (CMOD) was measured using in situ radiographic images. MATLAB was used to threshold and binarize the beam on either side of the notch, and the distance between the nearest points at the base of the resulting masks was computed to quantify CMOD. The maximum CMOD per cycle was then used to evaluate the rate of CMOD increase as a function of cycle number, $N$.

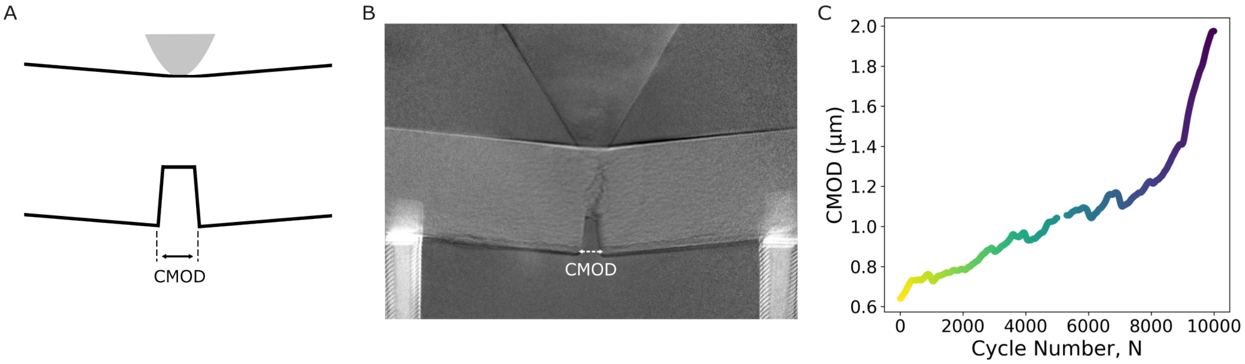


**Figure S4:** (A) Schematic showing CMOD as the displacement measured at the notch mouth. (B) Radiographic image from the experiment used for measuring CMOD; arrows indicate how we defined CMOD measurement. (C) Plot of CMOD vs. cycle number, highlighting the progressive increase in notch width opening throughout fatigue loading up to $10^4$ cycles.

#### Crack growth rate analysis of all samples

We analyzed crack growth rates across all samples, as shown in Fig. S5. Column A represents the specimen with a normalized stress intensity factor, $\tilde{K} = 0.15$, extensively characterized in the main text through fatigue loading analysis. Similar crack growth behavior, particularly deceleration patterns, were observed in Fig. S5Bi, corresponding to the specimen with $\tilde{K} = 0.25$. Both specimens displayed comparable initial crack growth rates, suggesting analogous decelerations at interfaces.

Figure S5, Column C represents the specimen exhibiting the highest initial crack growth rate, with $\tilde{K} = 0.3$ . This crack arrested within approximately 1 μm of crack growth due to deflection into a pre-existing defect as shown in the main text. Notably, Fig. S5Ai and Bi exhibited similar initial fatigue crack growth rates, whereas Fig. S5Ci shows distinctly elevated growth rates, as indicated by the different y-axis scale bar.

Rows ii, iii, and iv in Fig. S5 present the maximum applied load ($P_{max}$), crack extension ($\Delta a$), and crack growth rate ($da/dN$) respectively versus cycle number ($N$). Importantly, oscillations in $da/dN$ across all specimens, are independent of variations in the applied loading conditions.

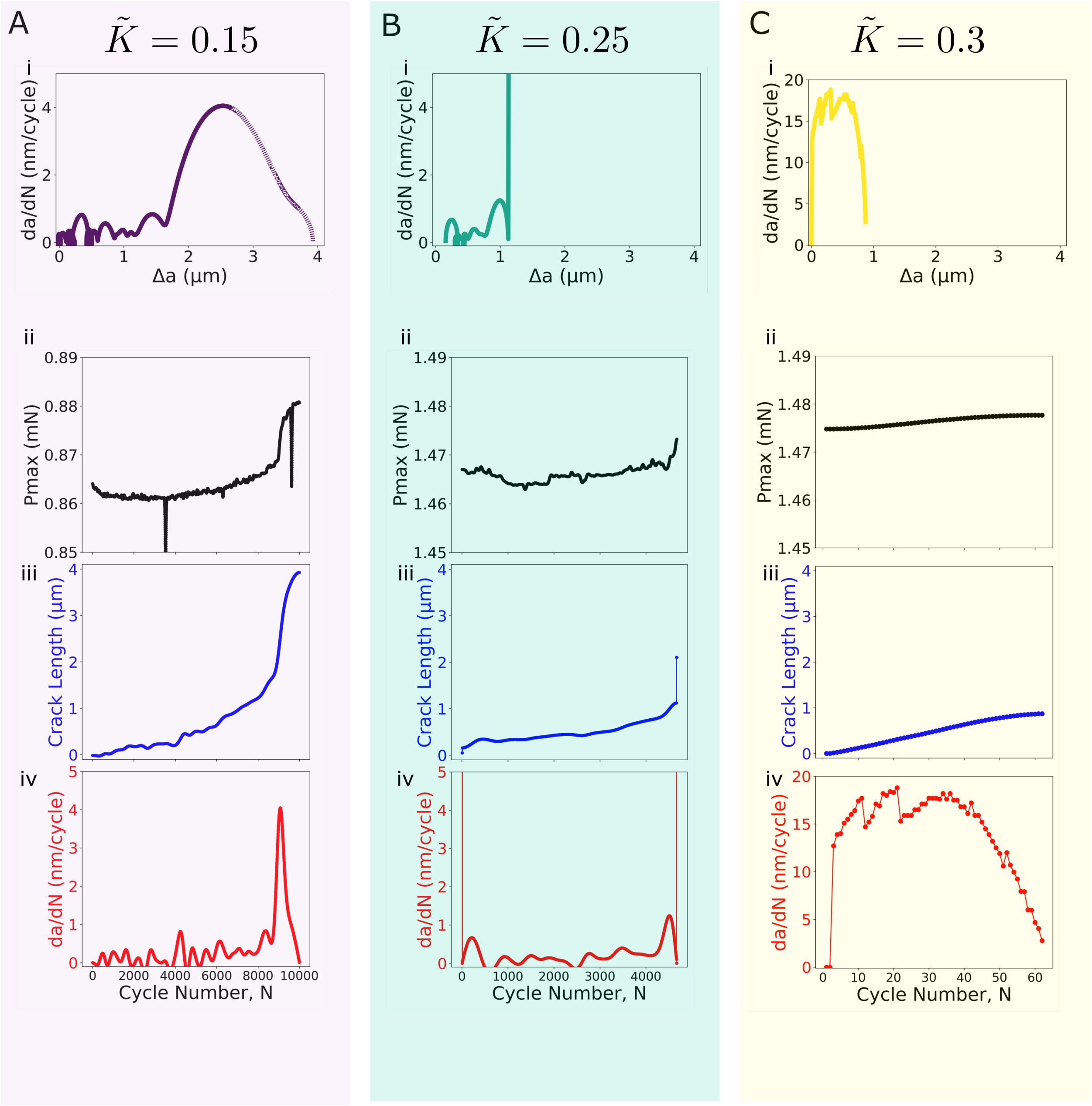


**Figure S5:** (i) Crack growth rate ($da/dN$) plotted against crack extension ($\Delta a$) for each specimen, highlighting deceleration patterns. Column A ($\tilde{K} = 0.15$), Column B ($\tilde{K} = 0.25$), Column C ($\tilde{K} = 0.3$, high initial crack growth rate with arrest). (ii) Maximum applied load ($P_{max}$), (iii) crack extension ($\Delta a$), (iv) and crack growth rate ($da/dN$) plotted against cycle number ($N$), highlighting load-independent decelerations in crack growth rates across specimens.

#### Processing and smoothing of mechanical data

The experimentally measured stiffness is smoothed with a Gaussian filter ($\sigma = 10$) as shown in Fig. S6A. This smoothed data is subsequently corrected as described in the previous section and then used to calculate crack length. Fig. S6B shows the smoothing of the calculated crack length using a Gaussian filter with $\sigma = 150$. This smoothed $\Delta a$ is used for subsequent stress intensity calculations. Fig.S6C shows the smoothing of the measured crack mouth opening displacement (CMOD) using a Gaussian filter with $\sigma = 150$. Finally, in Fig.S6D, both the smoothed $\Delta a$ and smoothed CMOD are plot, and this result is further smoothed using a Gaussian filter ($\sigma = 20$).

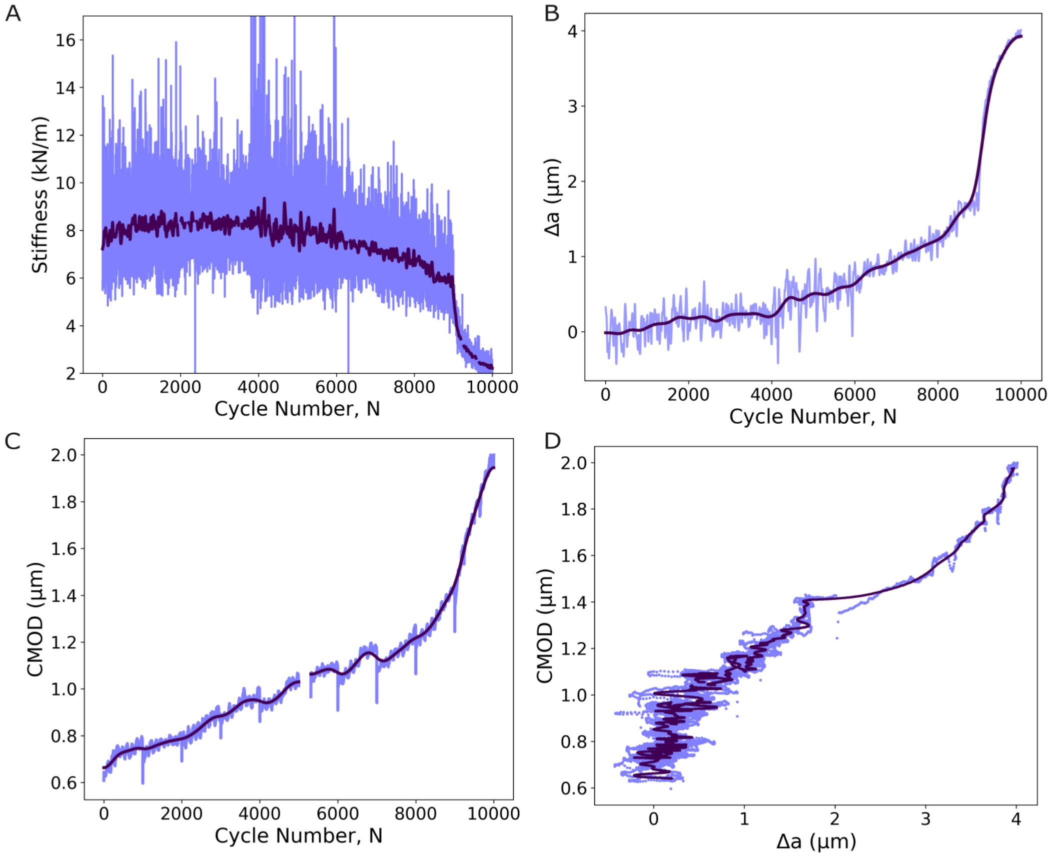


**Figure S6:** (A) Experimentally measured stiffness smoothed initially using a Gaussian filter ($\sigma = 10$), prior to correction and crack length calculation. (B) Calculated crack length data smoothed with a Gaussian filter ($\sigma = 150$), shown overlaying raw data. (C) Measured CMOD is smoothed with a Gaussian filter ($\sigma = 150$). (D) Smoothed CMOD vs smoothed $\Delta a$ are plot. This data is further smoothed with a $\sigma = 20$

### Sensitivity of Oscillation Widths to Smoothing

To evaluate whether the measured oscillation widths in the $\frac{da}{dN}$ vs. $da$ curve were sensitive to the level of Gaussian smoothing applied, we extracted peak widths from $\frac{da}{dN}$ computed with varying smoothing levels of $da$ ($\sigma$ = 100-200). For each smoothing level, peaks in the $\frac{da}{dN}$ curve were identified using consistent prominence and distance thresholds, and their widths were computed at half maximum height.

As shown in Fig. S7A, the $\frac{da}{dN}$ vs. $da$ profiles exhibit qualitatively similar oscillation structure across smoothing levels. To quantify these oscillations, we computed the width of each peak and visualized the resulting distributions across selected smoothing conditions (Fig. S7B. A one-way analysis of variance (ANOVA) revealed no statistically significant differences in peak widths across smoothing levels, F(5, N) = 0.585, $p$ = 0.675, indicating that the oscillation widths are not meaningfully influenced by the degree of smoothing within this range. This result suggests that the observed features in the $\frac{da}{dn}$ profiles are robust and not artifacts of the smoothing process.

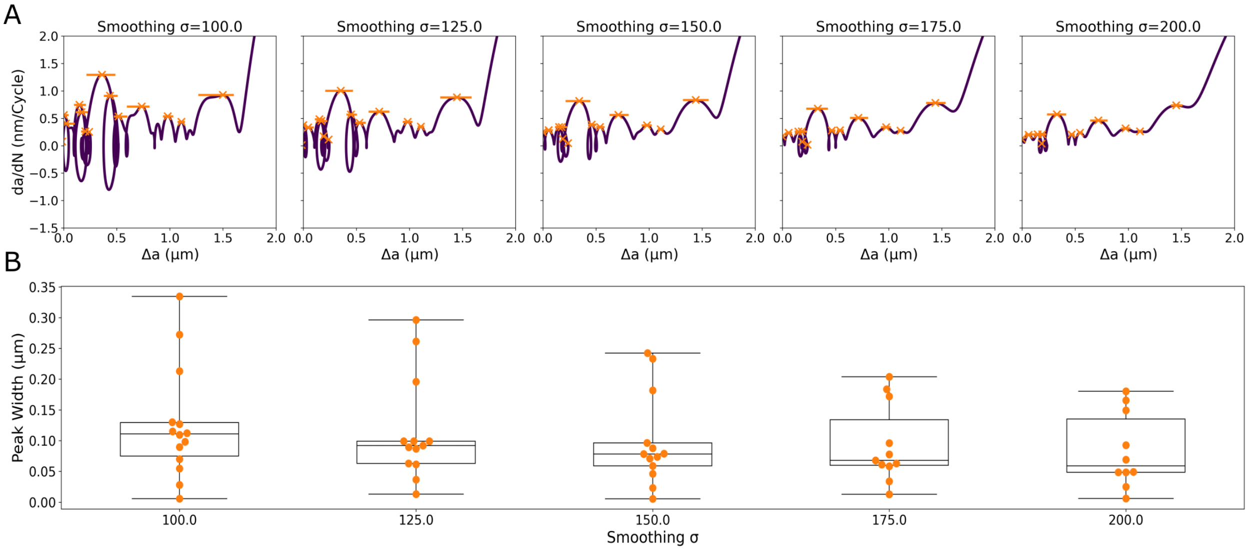


**Figure S7:** Sensitivity of oscillation widths in the $\frac{da}{dN}$ vs. $a$ profiles to Gaussian smoothing levels. (A) $\frac{da}{dN}$ vs. $a$ curves computed using different Gaussian smoothing values ($\sigma$ = 100–200), showing qualitatively similar oscillatory features across smoothing conditions. (B) Distribution of peak widths measured at half maximum height for each smoothing level. No significant differences were observed across smoothing levels, indicating that the oscillation widths are robust to the degree of smoothing applied.

#### Supplementary Videos

1. Fibril Bridging 2. Radiography loading experiment
